# Oligoribonuclease functions as a diribonucleotidase to bypass a key bottleneck in RNA degradation

**DOI:** 10.1101/546648

**Authors:** Soo-Kyoung Kim, Justin D. Lormand, Cordelia A. Weiss, Karin A. Eger, Husan Turdiev, Asan Turdiev, Wade C. Winkler, Holger Sondermann, Vincent T. Lee

**Affiliations:** Department of Cell Biology and Molecular Genetics, University of Maryland, College Park, MD, USA; Department of Molecular Medicine, College of Veterinary Medicine, Cornell University, Ithaca, NY 14853, USA

## Abstract

Degradation of RNA polymers is a multistep process catalyzed by specific subsets of RNases. In all cases, degradation is completed by exoribonucleases that recycle RNA fragments into nucleotide monophosphate. In γ-proteobacteria, a group of up to eight 3’-5’ exoribonucleases have been implicated in RNA degradation. Oligoribonuclease (Orn) is unique among them as its activity is required for clearing short RNA fragments, a function important for cellular fitness. However, the mechanistic basis for this substrate selectivity remained unclear. Here we show that Orn’s activity as a general exoribonuclease has been vastly overestimated by demonstrating that the enzyme exhibits a much narrower substrate preference for diribonucleotides. Co-crystal structures of Orn with substrates reveal an active site optimized for diribonucleotides that does not accommodate longer substrates. While other cellular RNases process oligoribonucleotides down to diribonucleotide entities, our functional studies demonstrate that Orn is the one and only diribonucleotidase that completes the final stage of the RNA degradation pathway. Together, these results indicate that Orn is a dedicated diribonucleotidase that clears the diribonucleotide pool that otherwise affects cellular physiology and viability.

## Introduction

Degradation of RNA is initiated by endonuclease-catalyzed cleavages; the resulting oligoribonucleotide fragments are hydrolyzed to completion by a mixture of exoribonucleases for the maintenance of cellular nucleotide pools^1^. Unlike the conserved machineries for the synthesis of macromolecules, distinct sets of RNases are used by different organisms to degrade these oligoribonucleotides. *E. coli* genomes encode eight 3’-5’ exoribonucleases, namely polynucleotide phosphorylase, RNase II, D, BN, T, PH, R and oligoribonuclease (Orn)^1,2^. A subset of these enzymes recognizes structural features of the RNA substrate, such as in tRNA, while others act on unstructured polymers^3^.

As an exoribonuclease, Orn is unique for two reasons. First, *orn* is required for viability in many γ-proteobacteria, including *E. coli*^4^ and other organisms^5,6^, unlike all other known 3’-5’ exoribonucleases. This indicates that not all exoribonucleases have redundant functions despite acting on the 3’ end of RNA substrates and in several cases, including RNase R and RNase II, sharing an activity toward oligoribonucleotides^7–10^. Hence, Orn appears to catalyze a particularly important step in RNA turnover. Second, Orn is a key enzyme in bacterial cyclic-di-GMP (c-di-GMP) signaling. The nucleotide second messenger c-di-GMP is produced in bacteria in response to environmental cues and controls a wide range of cellular pathways, including cell adhesion, biofilm formation, and virulence^11–14^. Since the discovery of c-di-GMP over 30 years ago, it has been known that the signal is degraded by a two-step process with a linear pGpG diribonucleotide intermediate^15^. While the enzyme for linearizing c-di-GMP to pGpG was discovered early on^16,17^, the identity of the enzyme that degrades pGpG remained elusive. Two recent studies showed that Orn is the primary enzyme that degrades pGpG in *Pseudomonas aeruginosa*^18,19^. In an *orn* deletion strain, the accrual of linear pGpG has a profound effect on cells. Specifically, the increase in pGpG inhibits upstream phosphodiesterases that degrade c-di-GMP, thereby triggering phenotypes associated with high cellular c-di-GMP levels. However, the molecular basis of Orn’s unique cellular functions in γ-proteobacteria that distinguishes it from all other exoribonucleases remains unexplained.

Since its discovery over 50 years ago^20^, Orn has been presumed to degrade oligoribonucleotides^21–23^. This notion largely derived from assays utilizing two types of substrates. In one series of experiments, ^3^H polyuridine (poly(U)) was incubated with Orn, other enzymes, or lysates and analyzed by paper chromatography, which offers limited resolution overall^21–23^. In a second set of experiments, Orn was incubated with oligoribonucleotides that had been tagged at their 5’ terminus by a large fluorophore. The products of this reaction were resolved by denaturing polyacrylamide gel electrophoresis, thereby allowing for detection of products with one or more nucleotides removed from the 3’ terminus^18,24–27^. In both instances, it was concluded that Orn could processively degrade “short” oligoribonucleotides. Yet it was not clear how Orn might selectively target “short” RNAs, rather than simply binding to the penultimate sequence at the 3’ termini of single-stranded RNA of any length.

Here, we sought to rigorously examine Orn’s substrate preferences, revealing its unique properties. To that end, we incubated Orn with 5’ ^32^P end-labeled RNAs of varying lengths and analyzed the products of that reaction over time. Our data show that Orn exhibits a surprisingly narrow substrate preference for diribonucleotides. This finding is in stark contrast to the previous studies that described a broader substrate length range and that originally gave oligoribonuclease its name. We sought to understand this remarkable substrate selectivity of Orn by determining the crystal structures of Orn in complex with pGpG and other linear diribonucleotides. These data reveal a structural basis for the diribonucleotide preference and identify key residues for recognizing diribonucleotides. Furthermore, we find that Orn is the only diribonucleotidase in *P. aeruginosa*. In addition, we show that other diribonucleotides, in addition to pGpG, can affect bacterial physiology. From this we propose a general model of RNA degradation, wherein a combination of exoribonucleases process oligoribonucleotides down to diribonucleotides and Orn completes RNA recycling by cleaving diribonucleotides to nucleoside monophosphates. In this way, Orn occupies an important but discrete step in the overall RNA degradation pathway.

## Results and Discussion

### Orn functions as a diribonucleotidase in vitro

To understand the length preference of Orn, recombinant affinity-tagged *Vibrio cholerae* Orn (Orn_*Vc*_) was purified and tested biochemically. We used an established ligand-binding assay^28^ to determine the relative substrate affinities of Orn to 5’-radiolabeled oligoribonucleotides that ranged from two to seven nucleotides in length. Unlike Cy-labeled RNA used in several earlier studies^18,24–27^, radiolabeling with ^32^P ensures that the substrate structure is unperturbed compared to native ligands. When spotted on nitrocellulose membranes, protein and protein-substrate complexes are sequestered at the point of application to the membrane, whereas unbound substrate diffuses due to radial capillary action^28^. By this assay, quantification of the diffusion zone reveals that Orn_*Vc*_ exhibits the highest affinity for diribonucleotide (*K*_*d*_ ^pGpG^ = 90 −/+ 9 nM), as compared to oligoribonucleotides of greater lengths (Table 1)^19^. Increase in the length to three or four residues reduces the affinity 7− or 10-fold, respectively. Substrates with 5 or more bases show a greater than 28-fold reduction in affinity compared to diribonucleotides. These results suggest that Orn has a strong preference for diribonucleotides over any other oligonucleotides.

**Table 1.**
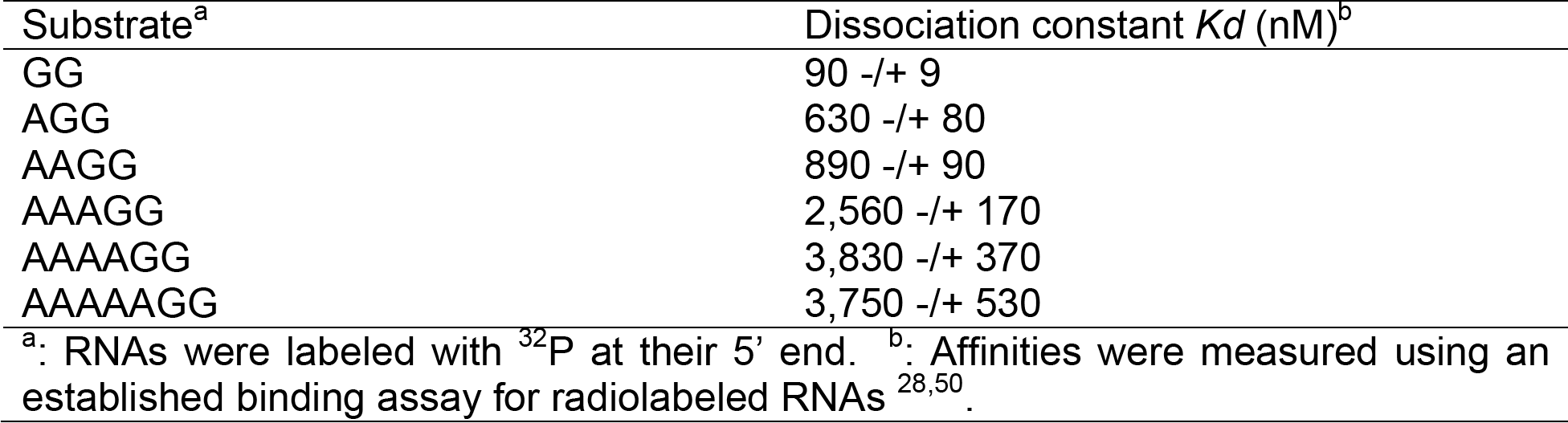
Quantitative measurement of length-dependent oligoribonucleotide affinities.

To understand whether the affinity preference reflects nuclease activity with natural substrates that are unmodified at the 5’ end, we incubated Orn_*Vc*_ with 5’-^32^P-radiolabeled oligoribonucleotides of varying lengths in the presence of divalent cations that support catalysis. The products of these reactions were resolved by urea-denaturing 20% polyacrylamide gel electrophoresis (PAGE). Under these electrophoresis conditions, the mononucleotides and oligonucleotides that were tested (between 2 and 7 nucleotides in length) can be resolved. The diribonucleotide substrate in this experiment was pGpG (GG), whereas the longer oligoribonucleotides included an increasing number of adenine nucleotides at the 5’ end. This arrangement ensured that the same GG sequence was maintained at the 3’ end while also avoiding stable G quadruplex formation that would be observed with RNAs containing stretches of poly-G^29^. At substrate concentrations that far exceed enzyme concentration (200:1) the diribonucleotide substrate was already fully processed to nucleoside monophosphates by 30 seconds (Figure 1A)^19^. In contrast, longer substrates (*i.e.*, from 3-mers to 7-mers) were not processed at their 3’ end, even at 30 minutes (Figures 1A and 1B). These results indicate that the length preference for exoribonuclease activity of Orn on diribonucleotides is even greater than mere differences in binding affinities. This strong substrate preference stands in stark contrast to previous studies arguing Orn acts as a general exoribonuclease that cleaves oligoribonucleotides with 2 to 7 residues in length^^18^,21,22,24,25^.

**Figure 1.**
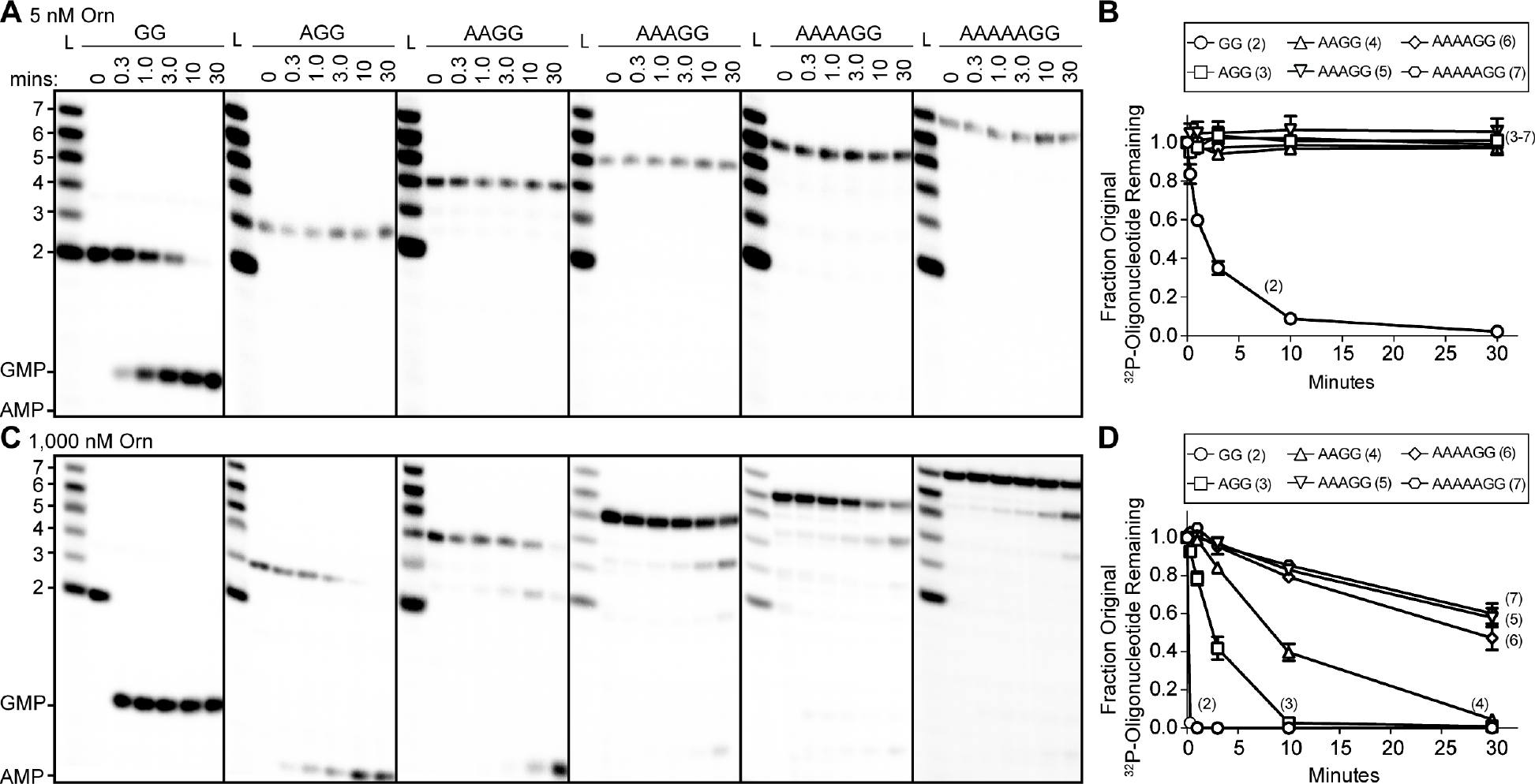
Orn has a stark preference for diribonucleotide cleavage *in vitro*. RNA nucleotides 2 to 7 residues in length (1 μM, containing the corresponding ^32^P-labeled RNA tracer) were each subjected to cleavage over time with 5 nM (A, B) or 1,000 nM Orn (C, D). Aliquots of each reaction were stopped at indicated times, and assessed by denaturing 20% polyacrylamide gel electrophoresis (A, C). Quantification of the intensities of bands corresponding to the amount of uncleaved initial oligonucleotide over time are plotted as the average and SD of three independent experiments (B, D).

To determine if Orn can indeed cleave substrates longer than a diribonucleotide, the enzyme was incubated with RNA substrates at a 1:1 molar ratio. Under these conditions, the diribonucleotide substrate was completely processed to nucleoside monophosphates by the earliest time point, 20 seconds (Figure 1C and 1D). Orn_*Vc*_ also facilitated the degradation of the longer RNA substrates, but only after significantly longer incubation times. For example, it required 10 minutes and 30 minutes to fully degrade 3-mer and 4-mer RNAs, respectively (Figures 1C and 1D). Cleavage was reduced further for longer RNAs; only 40-53% of the 5-mer, 6-mer and 7-mer RNAs were processed to nucleoside monophosphates at 30 minutes (Figures 1C and 1D), correlating with the weak affinities determined for these substrates (Table 1). Of note, we observed a non-uniform distribution of degradation products for the longer RNA substrates. Specifically, the diribonucleotide intermediate was never observed as a reaction intermediate for the longer RNAs. This reaction pattern indicates that RNAs of more than two residues could accumulate, but that diribonucleotide RNAs were always rapidly processed to nucleoside monophosphates. Together, these results indicate that Orn exhibits a strong substrate preference for diribonucleotides over longer oligoribonucleotides – far greater than reported previously^18,24^.

### Structure of Orn complexes with pGpG and other linear diribonucleotides

To elucidate the molecular basis for these unique properties of Orn, we set out to gain a deeper understanding of the enzyme’s substrate specificity by determining the crystal structures of two representative homologs, Orn_*Vc*_ ^6^ and the human REXO2 (also known as small fragment nuclease or Sfn^30^), bound to the diribonucleotide pGpG (Table S1; Figures 2A and 2B). Superimposing the two structures indicates their identical fold (rmsd of 0.75Å for the protomer; Figure S1A), which is preserved in the substrate-free Orn structure determined previously (rmsd of 0.54Å for the protomer; Figure S1A)^31^. The pGpG-bound structures reveal a narrow active site that is lined by the conserved acidic residues of the signature DEDD motif (D^12^, E^14^, and D^112^ of Orn_*Vc*_; D^15^, E^17^, and D^115^ of REXO2) and the general base H^158^ or His^162^ in Orn_*Vc*_ or REXO2, respectively (Figures 2A, 2B and S1B). In Orn_*Vc*_, the bases of the diribonucleotide buttress against aromatic residues W^61^ and Y^129^, the latter being contributed from the second half-side of the dimeric enzyme. The corresponding residues, W^64^ and Y^132^, are conserved in REXO2. Residue L^18^ in Orn_*Vc*_ (L^21^ in REXO2) wedges in-between the two bases. Most notably, residues S^108^, R^130^, S^135^ and the hydroxyl group on Y^129^ (S^111^, R^133^, S^138^, and Y^132^ in REXO2) form hydrogen bonds with the 5’ phosphate of pGpG, capping the substrate. This phosphate cap creates a major constriction of the active site, which is not observed in structurally related 3’-5’ exoribonucleases such as RNase T or ExoI (Figures 2C, S1C and S1D)^32–34^. RNase T and ExoI accommodate longer RNA substrates, facilitated by an expansive active site. The structural analysis correlates closely with sequence conservation of the phosphate cap motif, which is strict in Orn homologs but divergent in RNase T proteins (Figures 2D and S1B).

**Figure 2.**
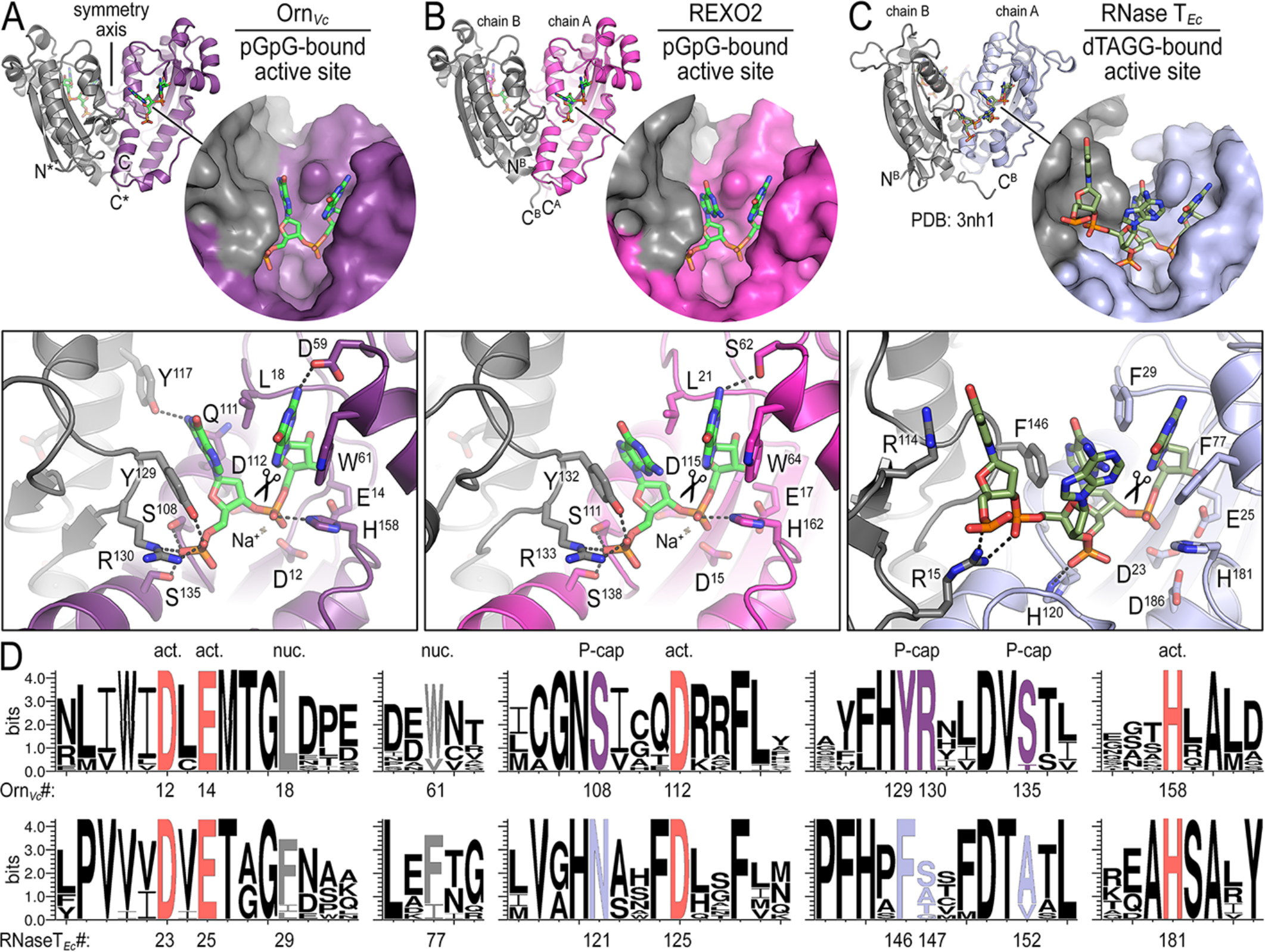
Structures reveal Orn’s conserved substrate preference for diribonucleotides. Crystal structures of pGpG-bound *V. cholerae* Orn (A) and human REXO2 (B) are shown in comparison to *E. coli* RNase T bound to substrate (PDB 3nh1; ^33^) (C), another DnaQ-fold 3’-5’ exoribonuclease with a DEDD(h) active site motif. The top panels show ribbon representations of the dimeric enzymes. The insets are surface representations of the enzymes’ active sites shown in similar orientations. The bottom panel describes the active site residues involved in RNA binding and catalysis. Residue numbering for REXO2 refers to its cytosolic isoform lacking the mitochondria-targeting pre-sequence. The sequence logos in (D) were constructed based on multi-sequence alignments of Orn and RNase T orthologs. Overall sequence identity ranges from 43-70% for Orn and 46-69% for RNase T. Sequence identifiers are provided in Table S2. Sequence logos were plotted using WebLogo ^51^. Conserved residues of the active site’s DEDD motif (‘cat.’; red), for ribonucleotide base binding (‘nuc.’; grey), and of the phosphate cap (‘P-cap’; purple) are highlighted.

### The phosphate cap is required for diribonucleotidase activity

To assess the impact on catalysis of the phosphate cap in comparison to other active site residues identified in the structural analysis, we introduced specific single-point mutations into Orn_*Vc*_. Purified proteins were evaluated for pGpG degradation. Mutations of the central phosphate cap residue R^130^ to alanine led to complete loss of catalytic activity, comparable to mutants in the DEDD active site motif (Figure 3A)^19^. We next asked whether Orn_*Vc*_ with specific point mutations could complement the deletion of *orn* in *P. aeruginosa*. *P. aeruginosa* Δ*orn* accumulates pGpG that in turn inhibits c-di-GMP-specific phosphodiesterases^18,19^. As a net result, c-di-GMP accumulates in these cells, an effect that is associated with a hyper-biofilm and cell aggregation phenotype. While expression of wild-type Orn_*Vc*_ complements the *P. aeruginosa* Δ*orn* resulting in a dispersed culture, complementation with variants that carry mutations in either the active site or phosphate cap residues results in cell aggregation indistinguishable from the Δ*orn* phenotype (Figure 3B). Together, these experiments demonstrate that an intact phosphate cap is required for enzyme function.

**Figure 3.**
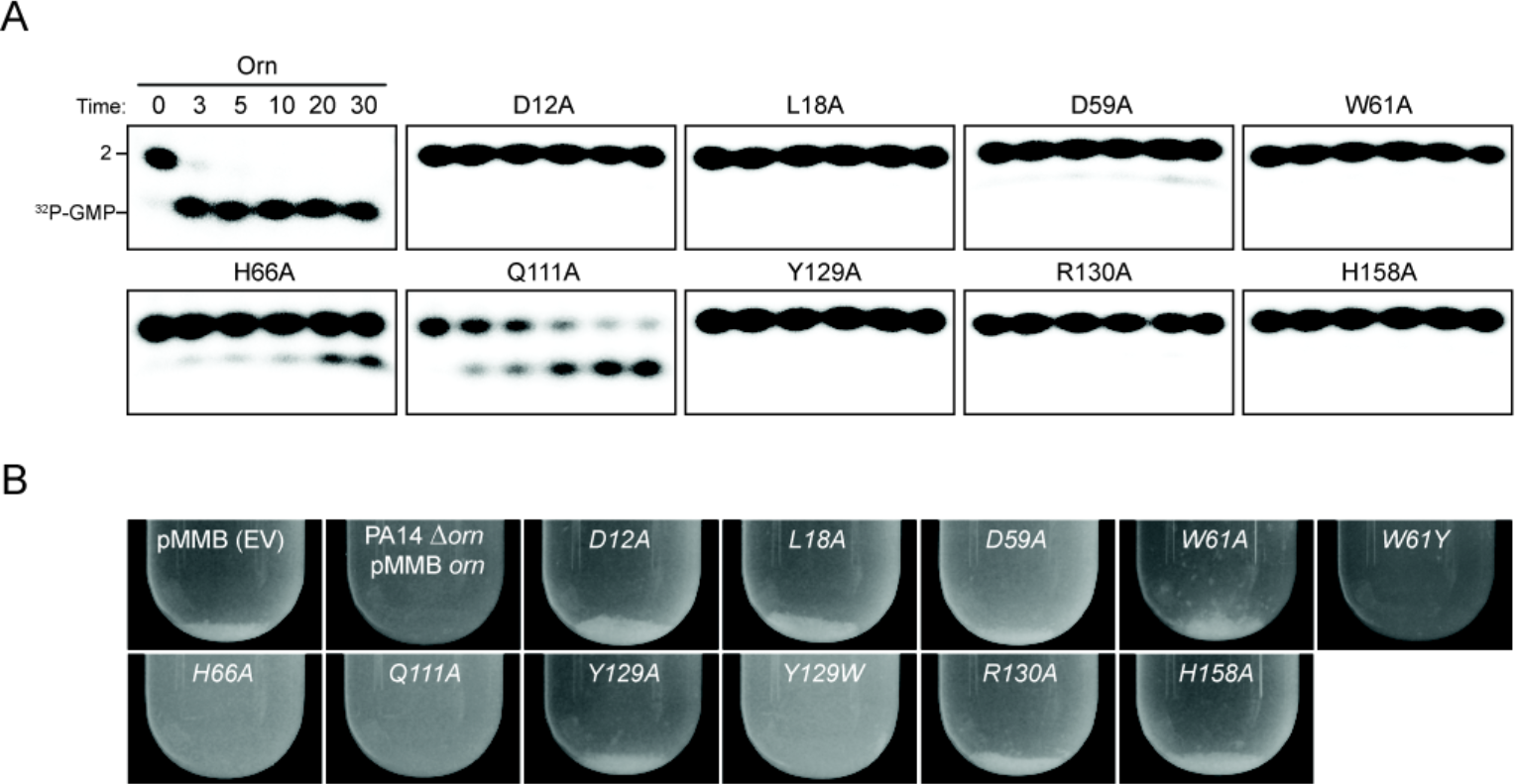
(Supplement to Figure 1A). Functional importance of active-site and phosphate-cap residues for Orn function. (A) *In vitro* enzyme activity. Degradation of ^32^P-pGpG (1 μM) by purified wild-type Orn_*Vc*_ or variants with alanine substitutions (5 nM) at the indicated sites was assessed. Samples were stopped at the indicated times and analyzed by 20% Urea PAGE. Representative gel images are shown with the indicated RNA size. The graphs show means and SD of three independent experiments. (B) *In vivo* activity of alanine substituted *orn*_*Vc*_ alleles to complement *P. aeruginosa Δorn*. Overnight cultures of the indicated strains were allowed to stand for 30 minutes without agitation to allow bacterial aggregates to sediment. Representative images of the cultures of triplicated assays are shown.

### Interaction of Orn with substrates

Structures of Orn_*Vc*_ (and REXO2) with different diribonucleotides, including di-purine, di-pyrimidine and mixed substrates, revealed identical binding poses (Figure S2). Additional hydrogen bonding between purine residues and Orn in the 3’ or 5’ position correlates with a small, but detectable preference of Orn_*Vc*_ for purine-containing substrates (Figure 4). Together, the structural analysis uncovered Orn’s mode of substrate binding, which is conserved from bacteria to humans and indicates a unique selection for linear diribonucleotides with a 5’ phosphate.

**Figure 4.**
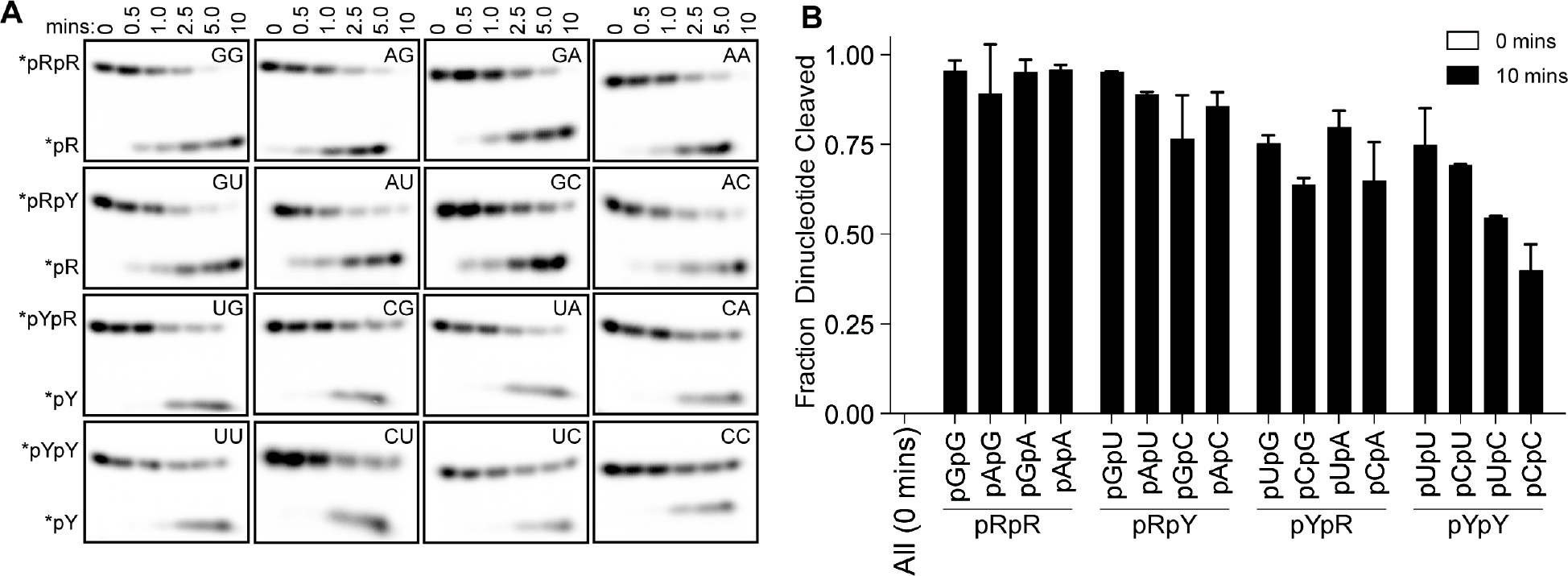
(Supplement to Figures 1 and S2B). Orn_*Vc*_ cleaves all diribonucleotides. (A) Orn_*Vc*_ (5nM) was incubated with di-purine (pRpR), purine-pyrimidine (pRpY or pYpR), or di-pyrimidine (pYpY) diribonucleotides (1 μM) containing the corresponding ^32^P-labeled RNA tracer. Aliquots of each reaction were stopped at the indicated times, and assessed by urea-denaturing 20% polyacrylamide gel electrophoresis. (B) Quantification of the intensities of bands corresponding to the amount of diribonucleotide cleaved at the 10 min timepoint. Results are the average and SD of duplicate independent experiments. Orn cleaves all diribonucleotides to nucleoside monosphosphates, albeit to varying extents. Diribonucleotides consisting of two purines (pRpR) were hydrolyzed most efficiently, with over 90% of starting RNAs processed by 10 minutes. A majority (>75%) of diribonucleotides with a 5’ purine (pRpY) were also processed by the 10-minute endpoint. However, diribonucleotides with a 5’ pyrimidines exhibited moderately reduced levels of cleavage. Di-pyrimidine (pYpY) substrates, and in particular pUpC and pCpC, showed the slowest turnover from the substrates tested. These results demonstrate that while all diribonucleotides are acceptable substrates for Orn, the enzyme is likely to exhibit moderate preferences for diribonucleotides that contain a 5’ purine.

Another distinguishing factor between Orn and other exoribonucleases with similar activities is the apparent inhibition of Orn by 3′-phosphoadenosine 5′-phosphate (pAp), a metabolic byproduct of sulfate assimilation that accumulates upon lithium poisoning of pAp-phosphatase^25^. DHH/DHHA-type oligoribonucleases such as NrnA from *Bacillus subtilis*, *Mycobacterium tuberculosis* and *Mycoplasma pneumoniae* are capable of dephosphorylating the pAp mononucleotide and their activity against oligoribonucleotides is unaffected by pAp^25,35^. In contrast, pAp has been described as a competitive inhibitor for *E. coli* Orn and the human REXO2^24^. However, the oligo-cytosine RNAs used in these prior studies were labeled at their 5’ end with a bulky fluorescent dye moiety (usually a cyanine or Cy-dye)^18,24^; our aggregate data now indicate that these RNAs represent suboptimal substrates for Orn, given its stark requirement for a simple 5’ phosphate cap (Figure 2). This labeling strategy is therefore likely to incompletely assess native Orn activity by underestimating diribonucleotidase activity while also overestimating the effect of competitive inhibitors. Revisiting pAp binding to Orn, we show here that Orn_*Vc*_ does not interact with radiolabeled pAp in the binding assay that was used to quantify oligoribonucleotide interactions with Orn (Figure S3). Furthermore, unlabeled pAp failed to competitively inhibit degradation of radiolabeled pGpG to GMP (Figure S3).

### Orn is the only diribonucleotidase in *P. aeruginosa*

All available literature assumes Orn is a 3’-5’ exoribonuclease that is responsible for processing of short (between 2-7) oligoribonucleotides. Yet our structural and biochemical data suggest that the enzyme exhibits such a striking preference for diribonucleotide substrates that the *in vivo* function of Orn as a general exoribonuclease should be reconsidered. Therefore, we developed experimental conditions to measure Orn’s activity in cellular extracts. The *orn* gene is required for viability in most γ-proteobacteria, including *Vibrio cholerae*; however, it is not essential for growth of *P. aeruginosa* under most conditions^18,19^. Therefore, lysates were generated from *P. aeruginosa* strains, including parental PA14, Δ*orn*, and Δ*orn* complemented with *orn_Vc_*. 5’-^32^P-radiolabeled 2-mer or 7-mer RNA was then added to each of these lysates (Figure 5). Aliquots of the mixtures were removed and analyzed by urea denaturing 20% PAGE at varying time intervals. Extracts from parental PA14 digested the entire radiolabeled diribonucleotide in less than 5 minutes (Figure 5A). In contrast, lysates from strains lacking Orn failed to show any signs of diribonucleotidase activity even at longer time points. Diribonucleotidase activity in the lysates could be restored by ectopic expression of *orn* from a self-replicating plasmid or by addition of purified Orn (Figure 5A). These data confirm that cellular Orn is required for degrading diribonucleotides and no other cellular RNase of *P. aeruginosa* can substitute for Orn activity.

When the ^32^P-7-mer RNA substrate was incubated in extracts from parental PA14 it was digested to a ladder of degradative intermediates including 6-mer, 5-mer, 4-mer and 3-mer RNAs (Figure 5B). Of note, while these intermediates and the final mononucleotide product accumulated over time, the 2-mer intermediate was never observed over the time course. In contrast, lysates from the Δ*orn* mutant specifically accrued the diribonucleotide intermediate with no apparent production of its mononucleotide products. Ectopic expression of plasmid-borne *orn* restored the degradation of the 7-mer to mononucleotides and the diribonucleotide intermediate could no longer be observed. Furthermore, the diribonucleotide intermediate that accumulated in the Δ*orn* lysate was fully processed upon addition of purified Orn protein (Figure 5B). Together, these results show that *P. aeruginosa* accumulates a bottleneck of diribonucleotide intermediates in a Δ*orn* background, which is only resolved upon addition of Orn. Degradation of RNA fragments with 3 or more residues by Orn is negligible in a cellular context, considering that Δ*orn* lysates preserve nuclease activities for the processing of RNAs down to diribonucleotides. From these aggregate data we propose that Orn functions not as an oligoribonuclease as stated in the literature but instead functions as a specialized ribonuclease of diribonucleotide substrates (*i.e.*, a ‘diribonucleotidase’).

**Figure 5.**
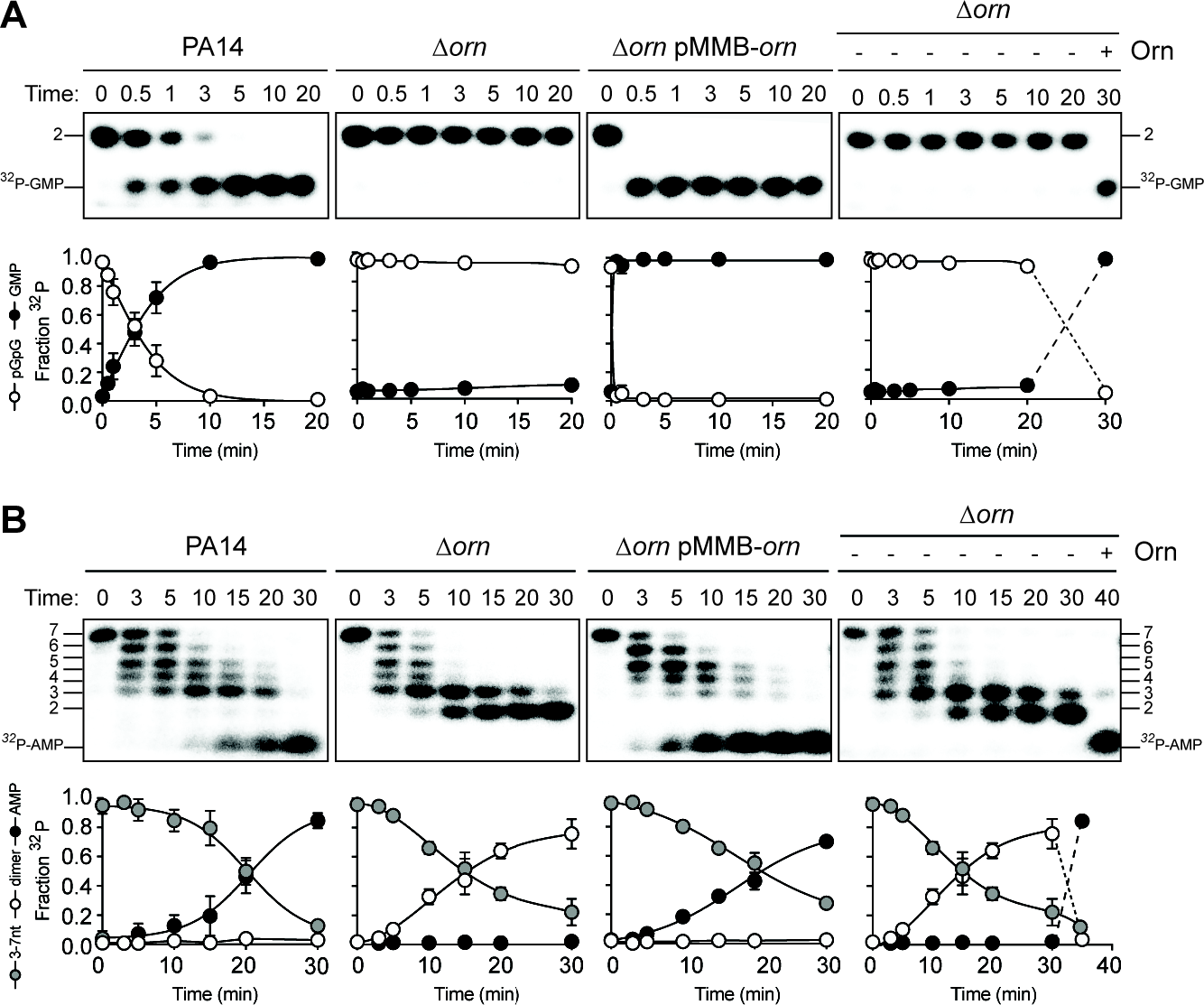
Orn acts as a diribonucleotidase in cell lysates. Degradation of ^32^P-GG (A) and ^32^P-AAAAAGG (B) by whole cell lysates of wild-type, *orn* mutant, *orn* mutant complemented with *orn*, or with 100 nM of purified Orn. Samples were stopped at the indicated time and analyzed by 20% Urea PAGE. Representative gel images of triplicated assays are shown with the indicated RNA size. Graphs show quantitation of triplicate data for indicated RNA species over time.

### The diribonucleotide pool affects *P. aeruginosa* growth in addition to c-di-GMP

We noticed that PA14 Δ*orn* had reduced growth on agar plates that is reminiscent of a small colony variant (SCV) phenotype (Figure 6)^36^. The SCV phenotype has been attributed previously to increased c-di-GMP levels^37^. C-di-GMP binds to FleQ and activates transcription of the *pel* operon^38,39^. In addition, c-di-GMP binds to the c-di-GMP-receptor PelD to increase the biosynthesis of the pel exopolysaccharide^40^, which enhances cell aggregation leading to a compact SCV morphology. Previous reports had shown that *P. aeruginosa* Δ*orn* is unable to clear pGpG, which results in the elevation of c-di-GMP signaling and pel-dependent cell aggregation and biofilm mass^18,19^.

**Figure 6.**
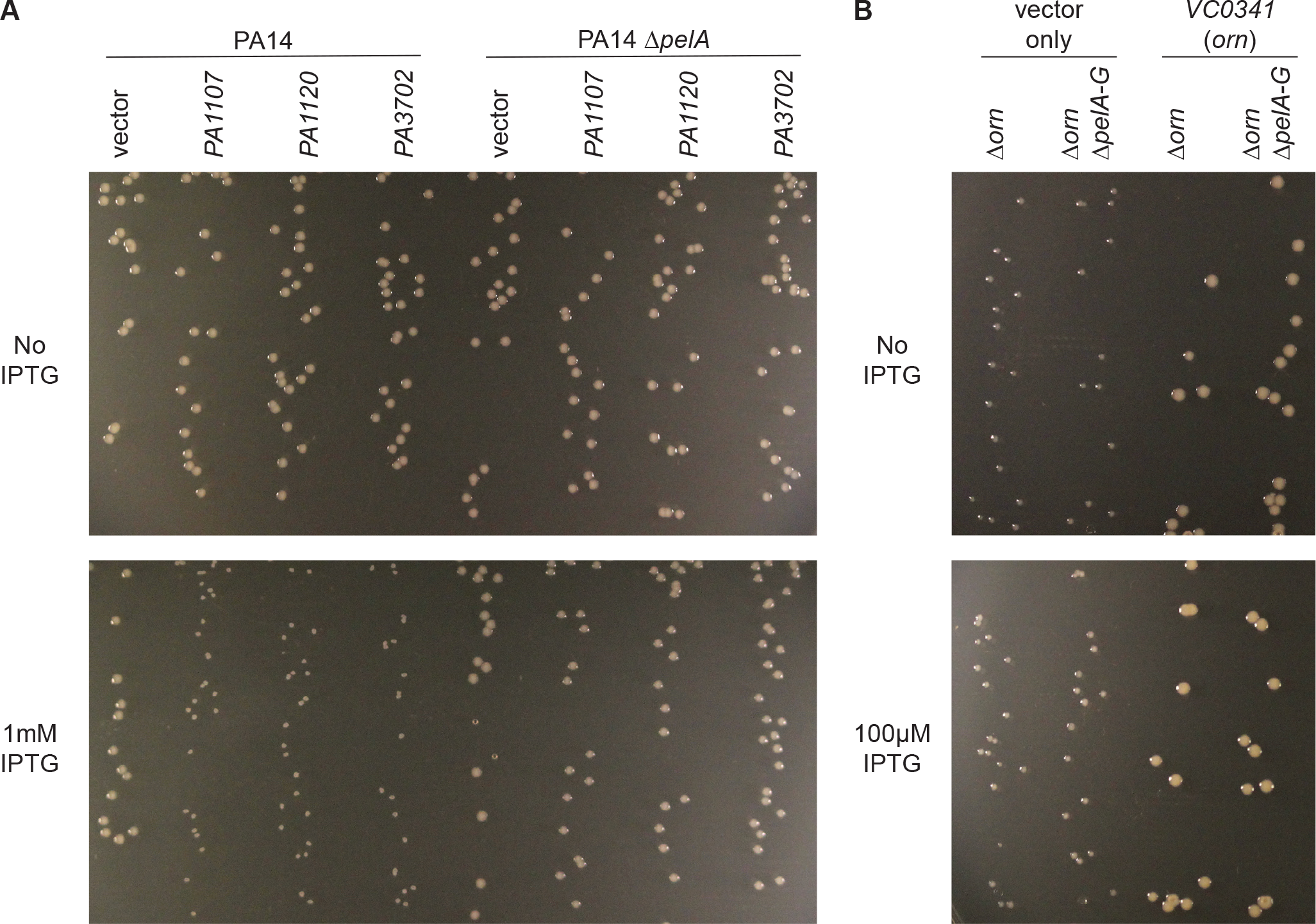
SCV phenotype of Δ *orn* is independent of c-di-GMP signaling. Bacterial cultures were diluted, 10 μL was spotted on LB agar plates with the indicated concentration of IPTG and drip to form parallel lines of bacteria. After overnight incubation, representative images of the plates were taken of the plates are shown for (A) PA14 and PA14 Δ*pelA* harboring pMMB vector with the indicated diguanylate cyclase gene and (B) PA14 Δ*orn* and PA14 Δ*orn* Δ*pelA-G* with pMMB and pMMB-*orn*. Experiments were performed in triplicate.

To determine whether the functional impact of diribonucleotide build-up is due specifically to an increased pGpG level and its effect on c-di-GMP, we asked whether the SCV formation in the Δ*orn* mutant is dependent on c-di-GMP processes. First, we confirmed that SCV formation in PA14 can be induced by elevated c-di-GMP levels through overexpression of diguanylate cyclases, such as PA1107, PA1120 and WspR (Figure 6A)^37,41–43^. As expected, the effect of c-di-GMP is due to increased production of the pel exopolysaccharide and biofilm formation since the PA14 Δ*pelA* mutant maintained normal colony morphology even when these diguanylate cyclases were overexpressed (Figure 6A). To determine whether the increase in the pool of pGpG and c-di-GMP is the only reason for small-colony growth in the Δ*orn* mutant, a Δ*orn* Δ*pel* double mutant was tested. When Δ*orn* Δ*pel* was grown on agar, it also had an apparent SCV phenotype (Figure 6B). Complementation of Δ*orn* and Δ*orn* Δ*pel* with active alleles of *orn* restored normal colony morphology, whereas inactive *orn* alleles failed to complement (Figure S4). In every case, the colony morphology was the same between Δ*orn* and Δ*orn* Δ*pel* indicating a second pathway that can restrict colony growth, but in this case independent of pel and c-di-GMP. These results indicate that increased pools of one or more of the diribonucleotides function to cause small-colony growth in Δ*orn* in addition to the actions of pGpG on c-di-GMP signaling.

## Conclusion

Orn is unique amongst exoribonucleases because it is essential in some γ-proteobacteria and is required to degrade the pGpG intermediate in c-di-GMP signaling. Yet the molecular and structural requirements for Orn were unknown. Our studies reveal that Orn is a dedicated diribonucleotidase in cells. This appears to be driven by a catalytic site that is restricted by a cap that mediates multiple interactions with the 5’ phosphate of diribonucleotide substrates. This constriction prevents longer substrates from binding with high affinity, rendering them poor substrates for catalytic cleavage. The discovery that Orn acts as a dedicated diribonucleotidase indicates that this activity clears a specific diribonucleotide bottleneck in global RNA degradation (Figure 7). Prior studies have shown that *orn* depletion can lead to accumulation of diribonucleotides and longer oligoribonucleotides in some cellular backgrounds^4^. The increase in oligoribonucleotides longer than dimers in cells may occur through feedback inhibition of other enzymes in the RNA degradation pathway, in analogy to the impact of pGpG on c-di-GMP-degrading phosphodiesterases and c-di-GMP signaling^18,19^. Whether the effect of diribonucleotides on essentiality is due to general processes such as RNA degradation, altered transcription^44^, specific interactions with essential proteins, or a combination of these remains to be evaluated in this context. Our studies therefore reveal a key step in RNA degradation, the enzymatic cleavage of diribonucleotides into mononucleotides, and set the stage to address how diribonucleotide accumulation is detrimental to cell survival.

**Figure 7.**
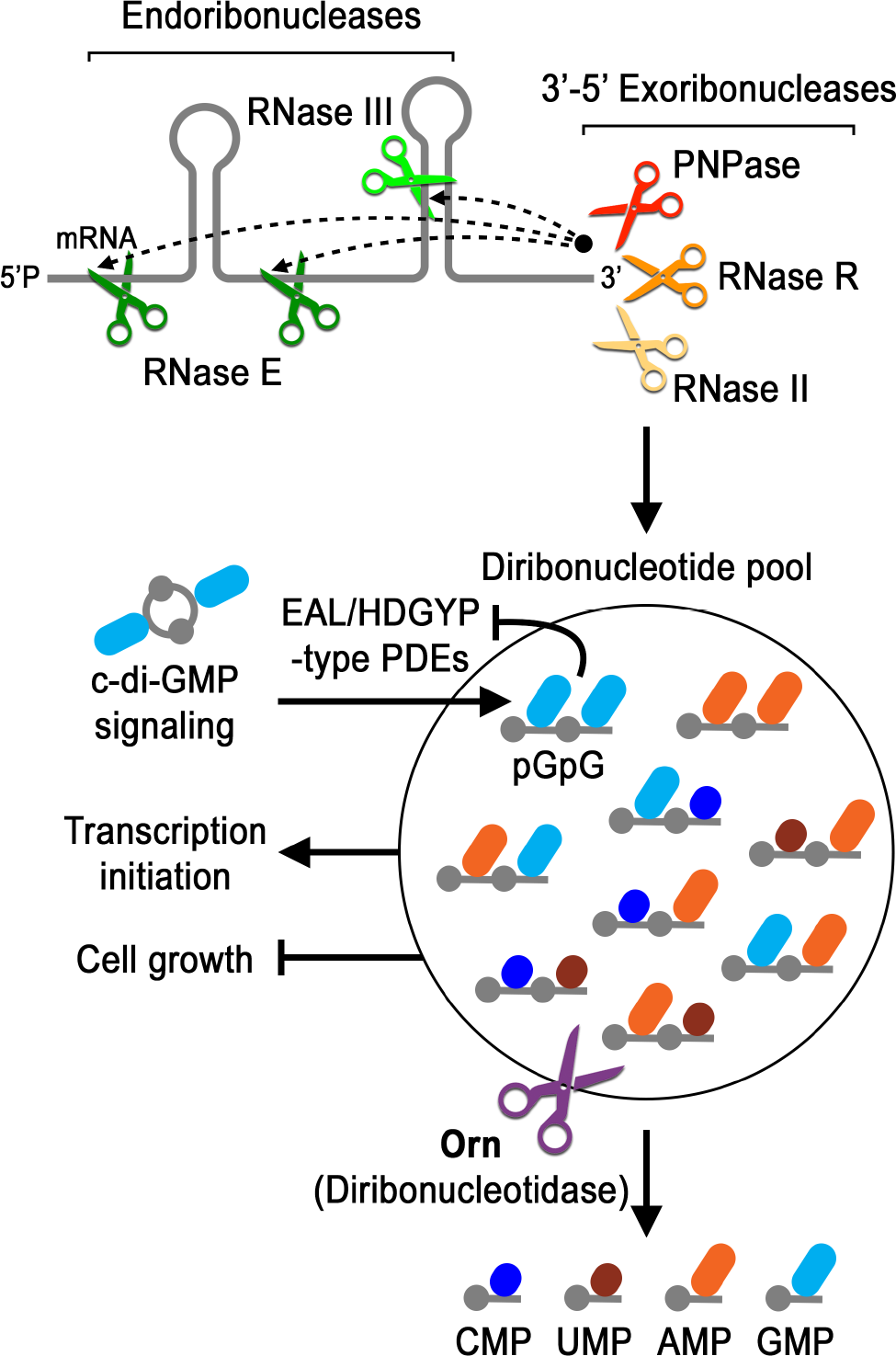
Model for Orn’s cellular function as a diribonucleotidase. RNA degradation is initiated by fragmentation via endoribonucleases (e.g. RNase E and RNase III) that cleave unstructured or structured RNA sequences. RNA fragments are processed further at their 3’ termini by 3’-5’ exoribonucleases (e.g. PNPase, RNase R, and RNase II). Their combined activity produces the various diribonucleotides from the original RNA substrate. The pGpG (GG) linear diribonucleotide is also produced by linearization of c-di-GMP by specific phosphodiesterases, EAL− and HD-GYP-domain-containing enzymes, which terminate c-di-GMP signaling. In *Pseudomonas aeruginosa* and likely other organism that rely on Orn for growth, Orn is the only diribonucleotidase that cleaves diribonucleotides to mononucleotides. In the absence of Orn, diribonucleotides accumulate with a drastic impact on cellular physiology, ranging from transcriptional changes, small-colony phenotypes and growth arrest, depending on the organism. Orn is also unique because it acts as the second phosphodiesterase in the degradation of c-di-GMP by clearing the pGpG intermediate. In an *orn* mutant, c-di-GMP levels are elevated through feedback inhibition of the c-di-GMP-degrading phosphodiesterases by pGpG, leading to the associated biofilm phenotypes.

## Methods

### Cloning, protein expression and purification

Orn genes from *Vibrio cholerae* O1 El Tor VC0341 (residues 1-181) and *Homo sapiens* REXO2 (residues 33-237) were synthesized by Geneart (Life Technologies). Genes were cloned by ligation between BamHI and NotI sites of a modified pET28a vector (Novagen) yielding N-terminally His_6_-tagged small ubiquitin-like modifier (SUMO) fusion proteins cleavable by recombinant Ulp-1 protease.

Orn proteins were overexpressed in *E. coli* BL21 T7 Express cells (New England Biolabs). Fresh transformants were grown in Terrific Broth (TB) supplemented with 50 ug/mL kanamycin at 37°C to an OD_600_ ~ 1.0, at which point the temperature was reduced to 18°C and expression was induced by addition of 0.5 mM IPTG. Cells were harvested after 16 hr of expression by centrifugation and resuspended in a minimal volume of Ni-NTA binding buffer (25 mM Tris-Cl, 500 mM NaCl, 20 mM imidazole, pH 8.5) followed by flash freezing in liquid nitrogen.

Cells were thawed and lysed by sonication. Cell debris was removed by centrifugation and clarified soluble lysate was incubated with Ni-NTA resin (Qiagen) pre-equilibrated with Ni-NTA binding buffer. Following one hour of binding, the resin was washed three times with 10 column volumes of Ni-NTA binding buffer, and then eluted with six column volumes of Ni-NTA elution buffer (25 mM Tris-Cl, 500 mM NaCl, 350 mM imidazole, pH 8.5). Eluates were buffer exchanged into gel filtration buffer (25 mM Tris-Cl, 150 mM NaCl, pH 7.5) via HiPrep 26/10 desalting column (GE Healthcare), followed by overnight incubation with Ulp-1 to cleave off the His_6_-tagged SUMO moiety. Untagged Orn proteins were recovered in the flow-through of a HisTrap Ni-NTA column (GE Healthcare), separated from His_6_-SUMO, uncleaved proteins and His_6_-tagged Ulp-1. Orn was concentrated by Amicon Ultra 10K concentrator prior to loading onto a HiLoad 16/60 Superdex 200 gel filtration column (GE Healthcare) equilibrated in gel filtration buffer. Fractions containing Orn were pooled and concentrated to 100 mg/mL, frozen in liquid nitrogen, and stored at −80°C.

### Protein Crystallography

REXO2-RNA and Orn_*Vc*_-RNA complexes (pGpG and pApA from BioLog Life Science Institute, other nucleotides from GE Healthcare Dharmacon) were formed prior to crystallization by mixing a 1:2 molar ratio of Orn:RNA in gel filtration buffer, followed by incubation for 30 min at the crystallization temperature. Orn-RNA complexes (10-30 mg/ml) were crystallized via hanging-drop vapor diffusion by mixing equal volumes (0.8 μl) of sample with reservoir solution. Orn_*Vc*_ crystals grew at 20°C over a reservoir solution that was composed of 0.1 M BisTris (pH 5.5), 17% polyethylene glycol 3350, and 20% xylitol. Crystals were flash-frozen in liquid nitrogen. REXO2 crystals grew at 4°C, using a reservoir comprised of 0.2 M sodium malonate (pH 5.5) and 15-20% polyethylene glycol 3350. REXO2 crystals were soaked in cryoprotectant of reservoir solution supplemented with 25% glycerol prior to flash freezing with liquid nitrogen. All crystals were stored in liquid nitrogen. Data were collected by synchrotron radiation at 0.977 Å on frozen crystals at 100 K at beamline F1 of the Cornell High Energy Synchrotron Source (CHESS). Diffraction data sets were processed using XDS, Pointless, and Scala ^45,46^. The initial structures were solved by Molecular Replacement using the software package Phenix ^47^ and the unpublished coordinates of *E. coli* Orn (PDB: 2igi) as the search model. Manual model building and refinement were carried out with Coot ^48^ and Phenix, respectively. Illustrations were prepared in Pymol (Version 2.2.0, Schrodinger, LLC). All software packages were accessed through SBGrid ^49^. Crystallographic statistics are shown in Table S1.

### Site-directed mutagenesis

To create the necessary mutants of *Vc0341 orn*, mutations were generated by using the Q5 Site-Directed Mutagenesis Kit (New England Biolabs).

### Labeling of RNAs

5’ un-phosphorylated RNAs were purchased from TriLink Biotechnologies or Sigma. Each RNA was subjected to radioactive end-labeling or non-radioactive phosphorylation by T4 Polynucleotide Kinase (New England Biolabs). Each RNA was subjected to phosphorylation with equimolar concentrations of either ^32^P-γ-ATP or ATP, T4 PNK, and 1X T4 PNK Reaction Buffer. Reactions comprising a final concentration of either 0.5 μM radiolabeled RNA or 2.0 μM phosphorylated RNA were incubated at 37°C for 40 minutes, followed by heat inactivation of T4 PNK at 65°C for 20 minutes.

### Protein expression and purification for biochemical assays

*E. coli* T7Iq strains harboring expression vector pVL847 expressing an His-MBP-Orn and His-MBP-Orn mutants from *V. cholerae* were grown overnight, subcultured in M9 fresh media supplemented with 15 μg/ml gentamicin and grown to approximately OD_600_ 0.5~1.0 at 30 °C. Expression was induced with 1 mM IPTG for 4 hours. Induced bacteria were collected by centrifugation and resuspended in 10 mM Tris, pH 8, 100 mM NaCl, and 25 mM imidazole. After addition of DNase, lysozyme, and PMSF, bacteria were lysed by sonication. Insoluble material was removed by centrifugation. The His-fusion protein was purified by separation over a Ni-NTA column. Purified proteins were dialyzed overnight against 10 mM Tris, pH 8, 100 mM NaCl, and 50% (vol/vol) glycerol, aliquoted, and frozen at −80 °C.

### Preparation of whole cell lysates

Overnight cultures of *P. aeruginosa* PA14 WT, Δ*orn* mutant, or complemented strains were subcultured into fresh media with antibiotic and 1mM IPTG, grown at 37°C with shaking. All bacteria samples were collected by centrifuge and resuspended in 100 mM KCl, 5 mM MgCl_2_, and 100 mM Tris, pH 8, also supplemented with lysozyme, DNase, and PMSF and stored at −80 °C.

### Oligoribonuclease cleavage reactions

Phosphorylated RNA (1.0 μM), including trace amounts of radiolabeled substrate, was subjected to cleavage by either 5.0 nM or 1.0 μM purified Orn at room temperature. These reactions were in the presence of 10 mM Tris-HCl pH 8.0, 100 mM NaCl, and 5 mM MgCl_2_. At the appropriate times, aliquots of the reaction were removed and quenched in the presence of 150 mM EDTA on ice. Aliquots were resolved by 20% urea-denaturing polyacrylamide gel electrophoresis (PAGE). The gel was imaged using Fujifilm FLA-7000 phosphorimager (GE). The intensity of the radiolabeled nucleotides was quantified using Fujifilm Multi Gauge software v3.0. Activity of whole cell lysates against ^32^P-labeled oligoribonucleotide substrates was measured by the appearance of truncated ^32^P-labeled products on 20% Urea PAGE containing 1× TBE buffer. The reactions were performed at room temperature in reaction buffer (10 mM Tris, pH 8, 100 mM NaCl, and 5 mM MgCl_2_). At indicated time, the reaction stopped by the addition of 0.2 M EDTA and heated at 98 °C for 5 min. Samples were separated on 20% urea-denaturing PAGE and analyzed as indicated above.

### DRaCALA measurement of dissociation constants

To measure K_D_, serial dilutions of purified His-MBP-Orn or His-MBP Orn mutants in binding buffer (10 mM Tris, pH 8, 100 mM NaCl, and 5 mM CaCl_2_) were mixed with radiolabeled nucleotides, applied to nitrocellulose sheets, dried, imaged and K_D_ values were calculated as described previously ^28,50^.

### Aggregation assay

A colony of each strain of *P. aeruginosa* grown in LB agar plates supplemented with 50 μg/mL carbenicilin was inoculated into borosilicate glass tubes containing 2.5 mL of LB supplemented with 0.1 mM IPTG. The cultures were placed in a fly-wheel in 37 °C incubator to spin for 18~22 hours. Culture tubes were allowed to settle at room temperature for 10 min and photographed.

### Data deposition

The atomic coordinates and structure factors have been deposited in the Protein Data Bank, http://www.rcsb.org (PDB ID codes 6N6A, 6N6C, 6N6D, 6N6E, 6N6F, 6N6G, 6N6H, 6N6I, 6N6J, and 6N6K).

## Acknowledgement

This work was supported by the NIH via grant R01GM123609 (to H.S.), R01AI110740 (V.T.L.), Cystic Fibrosis Foundation (CF Foundation) LEE16G0 (V.T.L.) and National Science Foundation (NSF) MCB1051440 (W.C.W.). C.A.W. was supported in part by the National Institutes of Health (NIH) training grant T32-GM080201. CHESS is supported by the NSF & NIH/NIGMS via NSF award DMR-1332208, and the MacCHESS resource is supported by NIH/NIGMS award GM103485.

---

**Figure S1.**
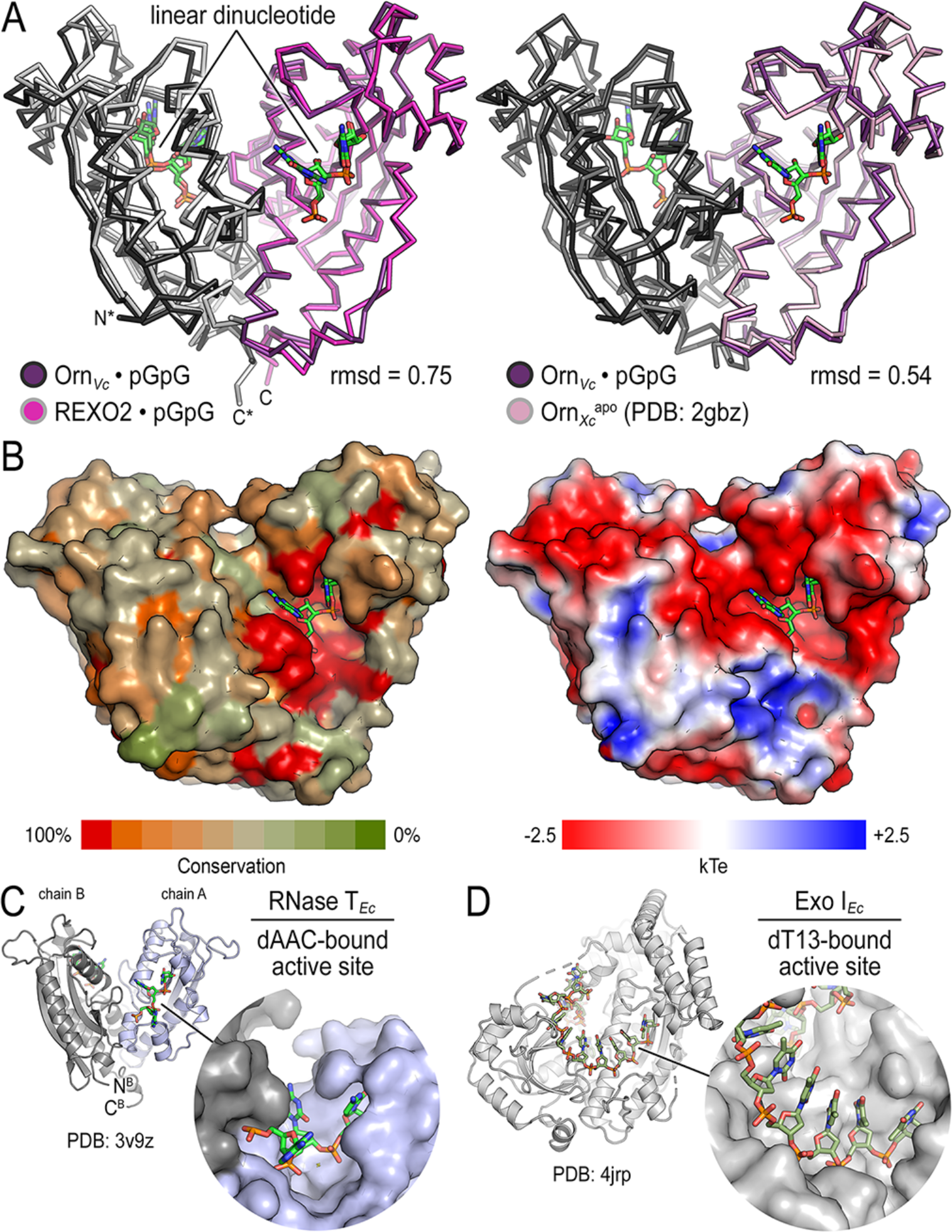
(Supplement to Figure 2). Structural comparison of Orn orthologs, RNase T and Exo I. (A) Superposition of Orn_*Vc*_ with REXO2 (left) and the substrate-free state of Orn from *Xanthomonas campestris* (PDB 2gbz; ^31^) (right). Rmsd values are reported for the alignment of a monomer. (B) Surface properties of Orn enzymes. An alignment of Orn orthologs was used to map conservation scores onto the solvent accessible surface of Orn_*Vc*_ (left). High degree of conservation overlaps with the acidic active site observed in the electrostatic potential map calculated with the APBS software ^52^ and based on Orn_*Vc*_ (right). (C) Active-site view of *E. coli* RNase T bound to dT13 (PDB 3v9z; ^32^). (D) Active-site view of *E. coli* Exo I bound to dAAC (PDB 4jrp; ^34^).

**Table S1.**
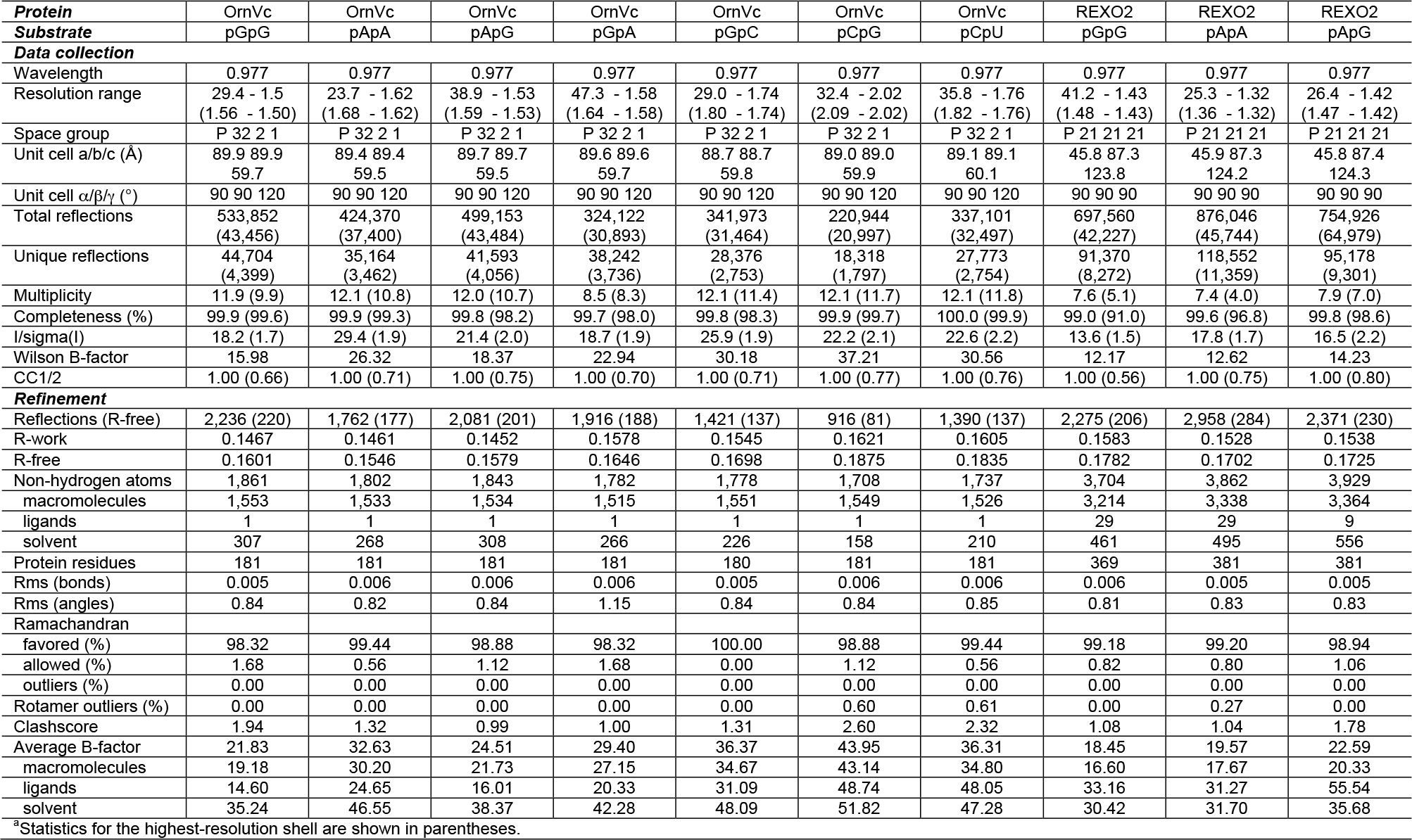
Data collection and refinement statistics.

**Table S2.** Sequences used to calculate surface conservation and generate Weblogos.

|  |  |  |  |  |  |
| --- | --- | --- | --- | --- | --- |
| Oligoribonuclease |  |  | RNase T |  |  |
| Strain | accession#<sup>a</sup> | %ID to *Orn<sub>vc</sub>*<sup>b</sup> | Strain | accession#<sup>a</sup> | %ID to *RNaseT<sub>vc</sub>*<sup>b</sup> |
| *Vibrio cholerae* | Q9KV17 | 100 | *Vibrio cholerae* | Q9KT97 | 100 |
| *Escherichia coli* | P0A784 | 68 | *Escherichia coli* | P30014 | 69 |
| *Coxiella burnetii* | Q83C93 | 59 | *Coxiella burnetii* | Q83B16 | 46 |
| *Haemophilus influenzae* | P45340 | 64 | *Haemophilus influenzae* | P44639 | 59 |
| *Acinetobacter baumannii* | V5VGJ9 | 59 | *Acinetobacter baumannii* | D0CDY9 | 53 |
| *Xanthomonas campestris* | Q8P8S1 | 55 | *Xanthomonas campestris* | Q8PAG3 | 47 |
| *Pseudomonas aeruginosa* | P57665 | 70 | *Pseudomonas aeruginosa* | Q9HY82 | 63 |
| *Pseudomonas fluorescens* | Q3KIZ7 | 69 | *Pseudomonas fluorescens* | Q3K7J7 | 61 |
| *Azotobacter vinelandii* | C1DLP5 | 66 | *Azotobacter vinelandii* | C1DQV0 | 65 |
| *Shigella flexneri* | Q83IJ9 | 67 | *Shigella flexneri* | Q0T4C2 | 69 |
| *Legionella pneumophila* | Q5ZRY0 | 62 | *Legionella pneumophila* | Q5ZRB8 | 52 |
| *Mycobacterium tuberculosis* | P9WIU1 | 43 |  |  |  |
| *Streptomyces coelicolor* | P57666 | 51 |  |  |  |
| *Homo sapiens* | Q9Y3B8 | 50 |  |  |  |

<sup>a</sup>: Based on Uniprot database (1); <sup>b</sup>: Based on ClustalOmega multi-sequence alignment scores (2).

**Figure S2.**
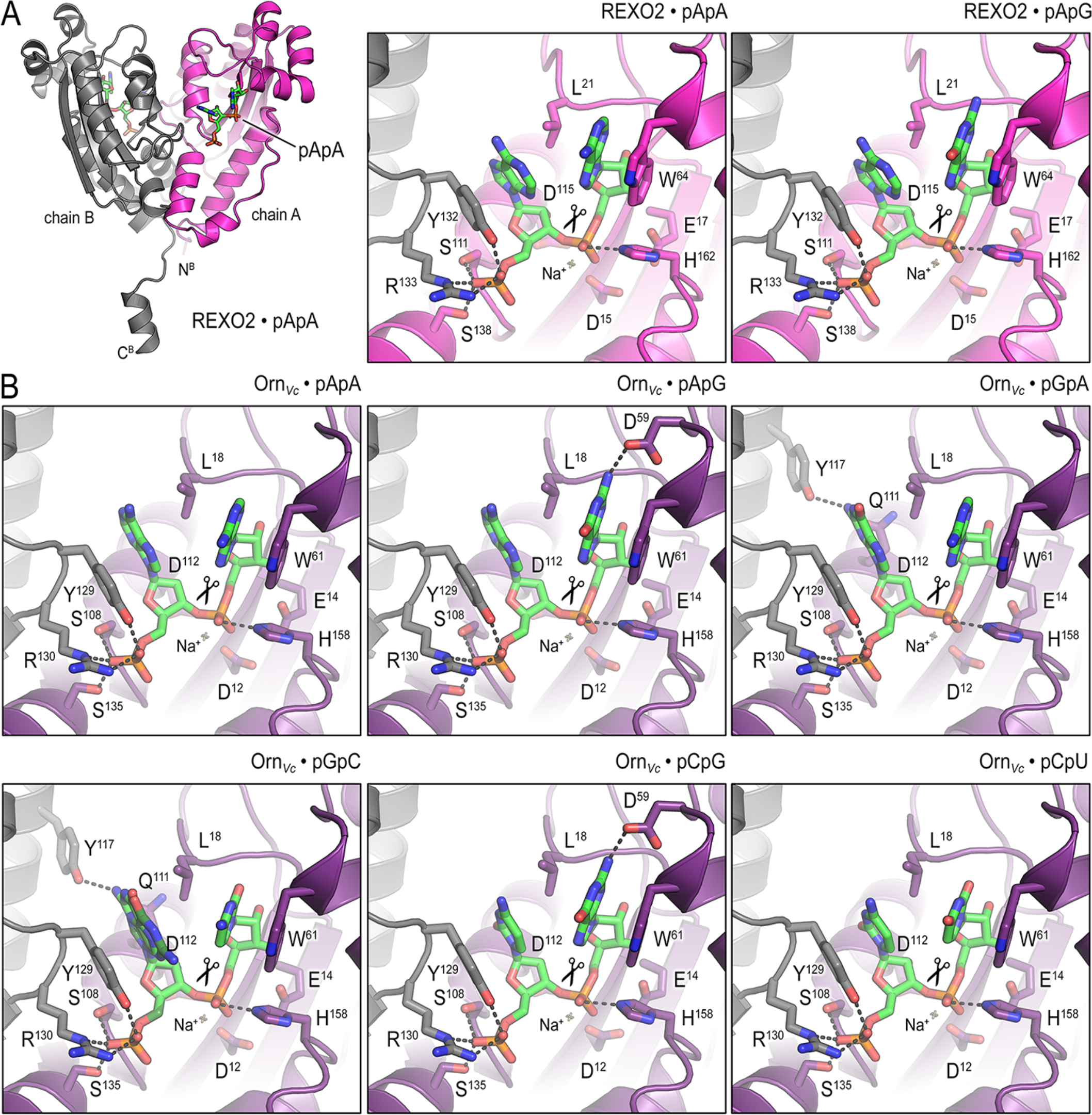
(Supplement to Figures 2A and B). Structural comparison of Orn _*Vc*_ and REXO2 bound to diverse diribonucleotides. (A) Crystal structures of REXO2 bound to pApA and pApG. The structural overview (left) shows a C-terminal helix that is unique to Orn orthologs in higher eukaryotes. (B) Crystal structures of Orn_*Vc*_ bound to di-purine (pApA, pApG, pGpA), purine-pyrimidine (pGpC, pCpG), and di-pyrimidine (pCpU) substrates. Views are identical to those shown in Figure 2 for the REXO2:pGpG and Orn:pGpG complexes.

**Figure S3.**
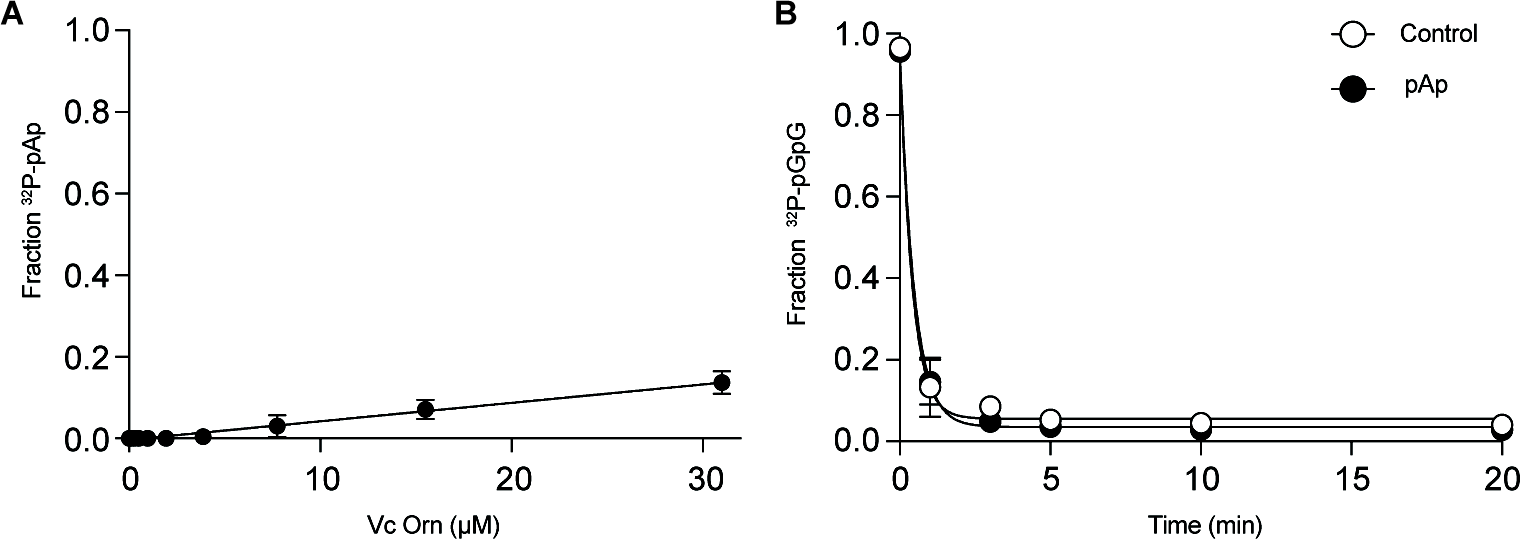
pAp has no effect on Orn activity. (A) The dissociation constant (K_d_) of ^32^P-pAp binding to Orn. K_d_ could not be measured as ^32^P-pAp binding to Orn was not saturatable. (B) The rate of ^32^P-pGpG degradation by Orn in the absence or presence of unlabeled pAp (100 mM). All data shown represent duplicate independent experiments.

**Figure S4.**
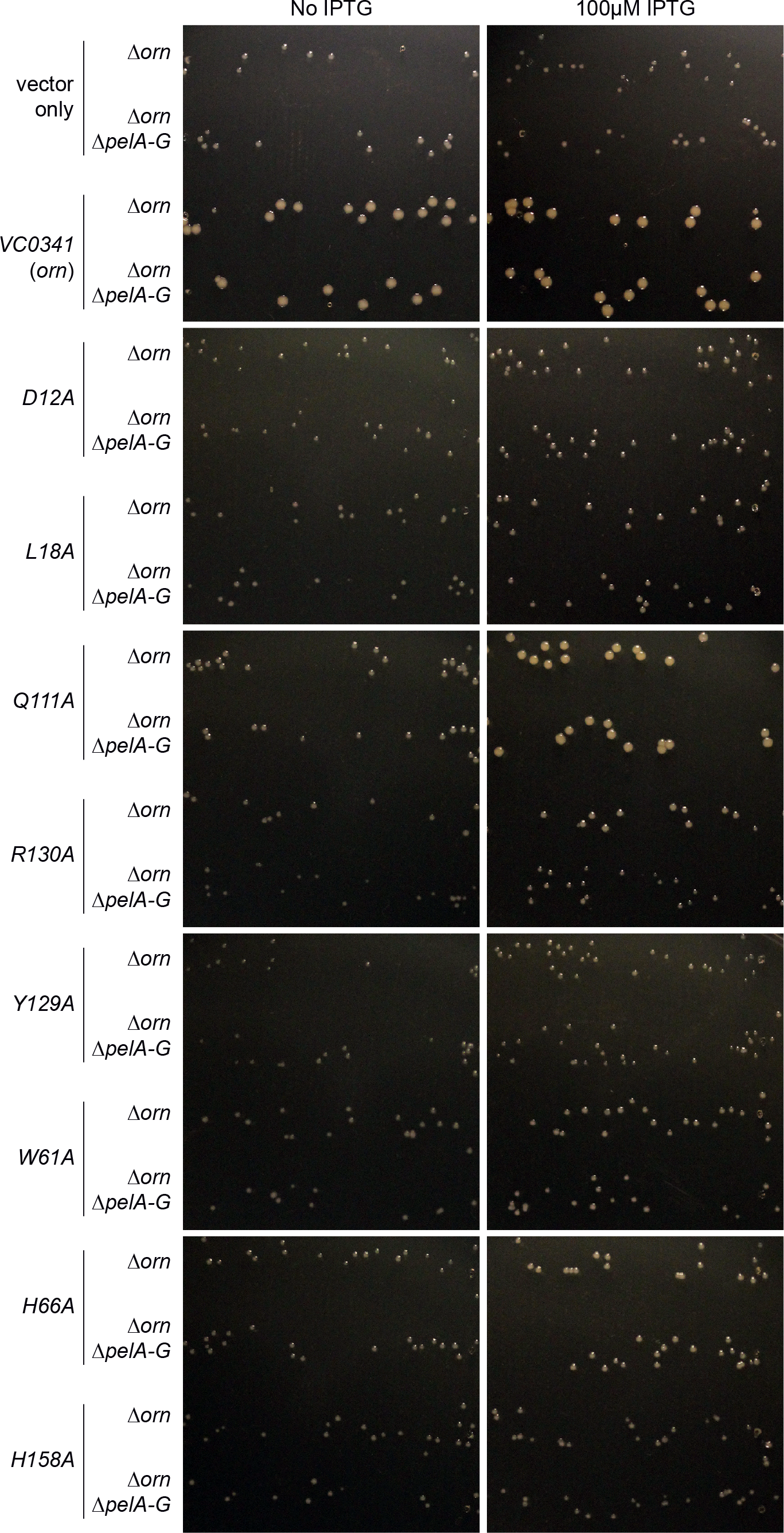
SCV phenotype of Δ *orn* is complemented by functional *orn* alleles. Bacterial cultures were diluted, 10 μL was spotted on LB agar plates with the indicated concentration of IPTG and drip to form parallel lines of bacteria. After overnight incubation, representative images of the plates were taken of PA14 Δ*orn* and PA14 Δ*orn* Δ*pelA-G* with pMMB alone or containing the indicated allele of *orn*. Experiments were performed in triplicate.

